# A Computational Model for Retinal Hemodynamics Under Gravitational and Postural Variations

**DOI:** 10.1101/2025.03.17.643768

**Authors:** Michele Nigro, Andrea Montanino, Eduardo Soudah

## Abstract

Understanding the effects of gravitational and postural variations on retinal circulation is crucial for both aerospace medicine and terrestrial health. This study presents a computational framework to analyze retinal hemodynamics under different gravitational conditions. A lumped-parameter model was developed to simulate blood flow, intraocular pressure, and vascular adaptations in response to changes in body posture and microgravity. Validation against experimental data demonstrated strong agreement in intraocular and ocular perfusion pressure variations across different tilt angles. Incorporating a simulated 4% constriction of the central retinal artery under microgravity conditions further model accuracy, highlighting the critical role of vascular remodeling. The simulation results indicate that posture significantly affects retinal circulation, leading to notable changes in vessel pressures, arterial velocities, and ocular perfusion. These findings emphasize the importance of fluid redistribution and vascular autoregulatory responses in conditions like Spaceflight-Associated Neuro-ocular Syndrome. This framework offers a powerful tool for developing countermeasures against vision-related risks in spaceflight and has potential applications in terrestrial ocular pathologies associated with altered intracranial pressures.

## Introduction

Spaceflight-Associated Neuro-Ocular Syndrome (SANS) is a major physiological challenge in human spaceflight [1]. SANS is characterized by a set of ocular and neuro-ophthalmic alterations observed in astronauts after prolonged exposure to microgravity, including optic disc edema (ODO), choroidal folds, globe flattening, and hyperopic shifts [2–6]. While initially attributed to intracranial pressure (ICP) elevation and alterations in the translaminar pressure gradient (TLP), recent evidence suggests that SANS is a multifactorial condition driven by cephalad fluid shifts, cerebral venous congestion, compartmentalized cerebrospinal fluid (CSF) dynamics, and impaired glymphatic clearance [5,7–10]. These interconnected pathophysiological mechanisms alter ocular perfusion, choroidal circulation, and retinal vascular autoregulation, leading to both structural and functional visual changes that may persist long after re-entry to Earth [11, 12]. Despite significant advances in space medicine and neuro-ophthalmic imaging, the precise pathophysiology of SANS remains incompletely understood [13–16]. Since in-flight ICP and ocular circulation measurements are very difficult to obtain [2, 17, 18], researchers rely on head-down tilt (HDT) bed rest studies as terrestrial analogs to simulate cephalad fluid shifts and investigate SANS-related mechanisms [11, 19–24]. However, to better quantify the complex interactions between gravitational changes, intracranial and ocular hemodynamics, computational models have been developed as complementary tools [25–30]. Computational models allow more detailed and refined exploration of ocular circulation under varying gravitational states [31–34]. Combining HDT experiments with numerical modeling has proven particularly effective for advancing our understanding of conditions such as SANS [33, 35]. Nevertheless, current ocular models still fail to capture the complexity and interactions between different ocular compartments fully. In particular, no model has yet been developed to investigate retinal circulation in relation to gravitational changes.

To address these limitations, we developed a computational model to simulate microgravity effects on retinal circulation by coupling fluid redistribution dynamics with variations in IOP due to changes in body inclination. Given its high metabolic demand and sensitivity to fluctuations in oxygenation and perfusion, the retina plays a central role in spaceflight-induced visual impairment [36–38]. As the neural interface between the eye and brain, its function depends on a delicate balance between intraocular and systemic circulatory factors [39, 40].

Recent findings by Iftime et al. [38] further highlight the susceptibility of the perivascular retina to gravitational shifts, showing that acute exposure to a microgravity analog significantly alters visual performance, particularly by increasing reaction times. These results emphasize the need for refined diagnostic approaches that extend beyond the *“central-fovea”* eye region to detect early manifestations of SANS.

By integrating these insights, our approach enabled analysis of key hemodynamic parameters (blood pressure distribution, flow velocity, and retinal blood inflow) critical to ocular health. The computational outcomes were validated against available experimental data, thereby enhancing our understanding of the ocular consequences of altered gravitational conditions.

Similarly, in Salerni et al. [32], a model consisting of the brain, eye, and aqueous humor circuit was presented, in which microgravity conditions were simulated by implementing an HDT and modifying the value of the filtration coefficient between the capillaries in the brain and the brain tissue, as well as the reflection coefficient. Conversely, in Fois et al. [33], along with a multiscale model of the cardiovascular system, an HDT maneuver was simulated alongside an experimental campaign, using the same model for IOP dynamics but without considering any interaction with the retinal vasculature.

A notable innovation of our model is its integration of two distinct yet interconnected models: a retinal vascular network model from Guidoboni et al. [41] and an IOP model responsive to gravitational orientation defined in Nelson et al. [31]. By coupling these components, the model provides a physiologically accurate representation of ocular circulation, reflecting realistic interactions between IOP fluctuations and retinal hemodynamics. In line with the approach originally taken by Guidoboni et al. [41] for glaucoma, but with a dynamic variation of IOP, arterial and venous blood pressures that depend on gravitational influence induced by changes in body inclination. This enhancement allows for a more comprehensive analysis of the interplay between posture-dependent IOP fluctuations and retinal circulation. This integrative approach is particularly relevant to SANS, as it facilitates detailed estimations of retinal blood flow, pressure distribution, and flow velocity under simulated microgravity conditions. Ultimately, this study offers a physiologically robust framework to investigate spaceflight-induced ocular alterations, supporting the development of targeted countermeasures to safeguard astronaut ocular health.

This paper is structured as follows. First, we introduce the computational model for retinal hemodynamics, providing an overview of its general framework before detailing the mathematical formulation of the ocular vascular network, the dynamic interaction between retinal circulation and IOP, and the effects of gravitational and postural changes. Next, we explain how the microgravity conditions are incorporated into the computational model, considering hydrostatic pressure redistribution and systemic circulatory adaptations. The results section then presents the simulated hemodynamic responses under different gravitational conditions, analyzing variations in blood pressure, retinal blood flow, and vascular autoregulation across multiple tilt angles, followed by a critical discussion of the results.

## Materials and methods

The ocular vascular system is represented through a numerical model shown in Figure 1. The retinal vasculature is divided into five main compartments: the central retinal artery (CRA), arterioles, capillaries, venules, and the central retinal vein (CRV). To describe the hemodynamics in these compartments, we employ a lumped modeling approach, which is widely used in cardiovascular studies due to its ability to capture complex interactions through a simplified circuit analogy [27, 42–45].

**Fig 1.**
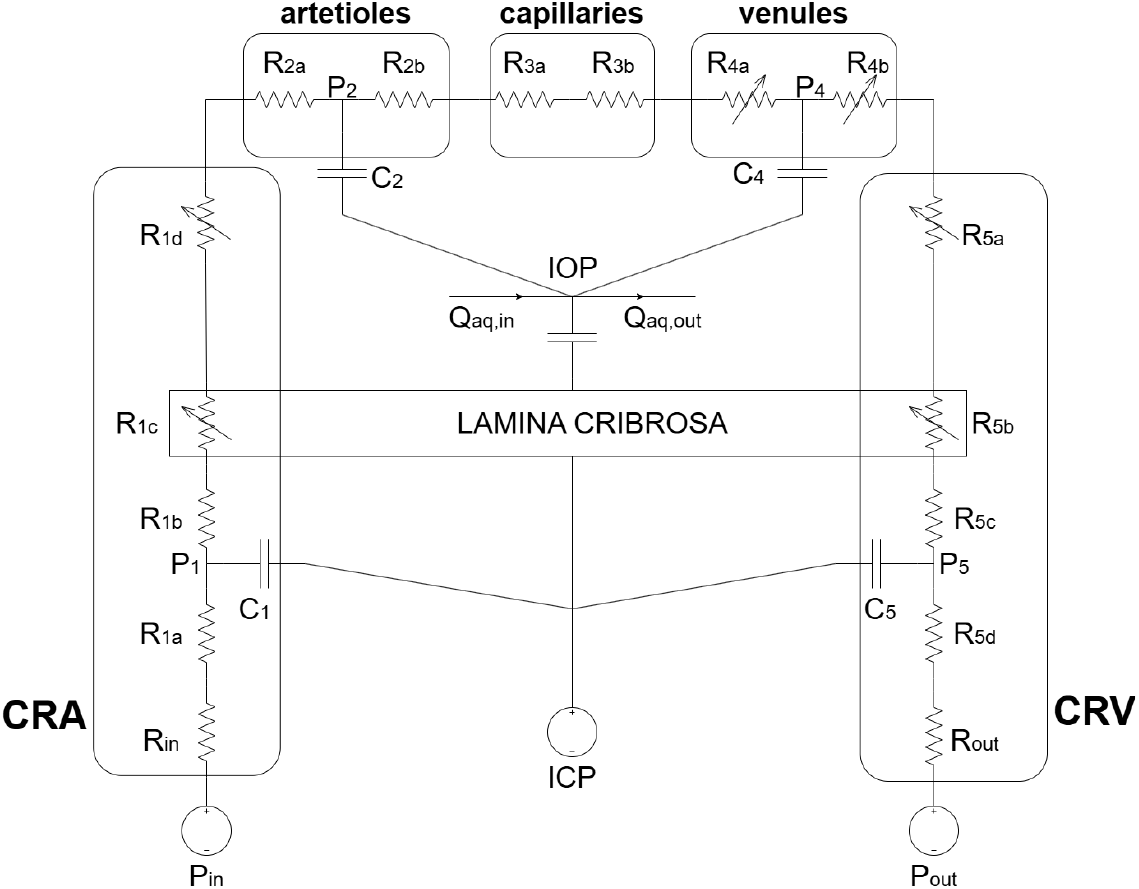
Lumped-parameter model of retinal hemodynamics. The circuit-based representation models the retinal vasculature using resistances (*R*) for vascular resistance and capacitances (*C*) for vascular compliance. The intraocular pressure (*IOP* ) dynamically interacts with aqueous humor inflow (*Q*_aq,in_) and outflow (*Q*_aq,out_). The lamina cribrosa acts as a transitional boundary, influenced by intraocular and intracranial pressures (*ICP* ). Input pressure (*P*_in_) and output pressure (*P*_out_) define the driving pressure gradient for retinal circulation. The model accounts for systemic and gravitational effects, allowing for the analysis of hemodynamic adaptations under altered postural conditions.

Within this computational framework, vascular segments are represented as elements of an electrical circuit, where pressure corresponds to voltage, flow to current, and vascular characteristics are modeled using resistors and capacitors. Moreover, lumped models are computationally efficient, striking a balance between accuracy and complexity compared to 1D and 3D models, while still providing meaningful insights [43, 46, 47]. For the equations related to compliance and vascular resistance, as well as other key hemodynamic relationships, see Table 1.

**Table 1.** Summary of Governing Equations in the Lumped Parameter Model.

|  | Equation |
| --- | --- |
| Mass Conservation | $Q = \frac{dV}{dt}$ |
| Ohm's Law in Hemodynamics | $\Delta P = QR$ |
| Poiseuille's Law (Resistance) | $R = \frac{128\mu L}{\pi D^4}$ |
| Compliance for a Cylindrical Vessel | $C = \frac{\pi D^3 L}{4Eh}$ |
| Pressure Evolution in a Compliant Vessel | $\frac{dP}{dt} = \frac{Q_{in} - Q_{out}}{C}$ |
| Blood Flow Rate (Poiseuille's Law) | $Q = \frac{\pi V D^2}{8}$ |
| Blood Velocity | $v(t) = \frac{8}{\pi D^2} \frac{\Delta P(t)}{R}$ |
| Resistivity Index (RI) | $RI = \frac{PSV - EDV}{PSV}$ |
| Mean Arterial Pressure (MAP) | $MAP = \frac{SBP + 2 \times DBP}{3}$ |
| Ocular Perfusion Pressure (OPP) | $OPP = MAP_{eye} - IOP$ |

Each vascular segment is exposed to different external pressures depending on its position within the network. Intraocular segments are subjected to IOP, retrobulbar segments experience the Retrolaminar Rissue Pressure (RLTp), which can be assumed to be equal to ICP [33], while translaminar segments are influenced by an external pressure that depends on the internal stress distribution within the lamina cribrosa. These pressure interactions play a crucial role in determining the mechanical and hemodynamic behavior of the ocular system.

Unlike previous models that considered IOP as a fixed boundary condition, our approach models the IOP as a dynamic state variable. This advance is based on the initial mathematical model proposed by Nelson et al. [31], and subsequently refined by Petersen et al. [35] and Fois et al. [33].

This approach enables a dynamic coupling between IOP and ocular hemodynamics, allowing the model to capture feedback mechanisms essential for understanding transient effects and autoregulatory responses under altered gravitational conditions, such as microgravity and head-down tilt maneuvers. By integrating the IOP variation intrinsically within the system, the model provides a biomechanically justified representation of ocular blood flow dynamics and their interaction with systemic circulatory changes.

The mathematical formulation of our model, including all governing equations, parameter definitions, and detailed derivations, is comprehensively presented in the Appendix and supplementary materials accompanying this article.

As the primary input parameter for the numerical model, the arterial blood pressure upstream of the eye was selected. In the lumped modeling framework, this pressure acts as the driving force for hemodynamic flow through the retinal vasculature, analogous to a voltage source in an electrical circuit. The relationship between pressure, flow, and resistance follows Ohm’s law in hemodynamics (Table 1).

Guidoboni et al. [41] estimated patient-specific values for the *P*_in_ and *P*_out_ profiles using an inverse problem approach. This estimation was based on blood flow velocity measurements acquired via bidirectional laser Doppler velocimetry in the CRA and CRV. These velocity profiles, when combined with the governing equations of the model, including Poiseuille’s law for resistance and vascular compliance equations (Table 1), allow for a physiologically accurate reconstruction of ocular hemodynamics, ensuring that the model captures the key pressure-flow interactions regulating retinal perfusion.

Regarding the *P*_out_, i.e., the venous pressure downstream of the eye, we adopted the condition proposed in the model by Petersen et al. [35]. To maintain physiological venous outflow, the model ensures that *P*_out_ remains above the central venous pressure (CVP), set at 7 mmHg. When episcleral venous pressure (EVP) is greater than CVP, *P*_out_ follows EVP; otherwise, it is equal to CVP. This relationship is expressed as a piecewise function:

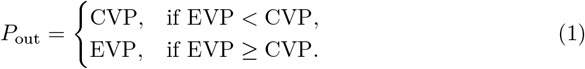

This formulation ensures that venous outflow remains physiologically viable under different gravitational conditions, as EVP varies with tilt angle and systemic circulatory changes. The dynamic interaction between EVP and CVP is particularly relevant in conditions such as microgravity and HDT, where venous return and episcleral venous drainage are altered.

To simulate the HDT maneuver, we considered multiple inclination angles relative to the horizontal axis. The *P*_in_ that in our model was assumed to correspond to a standing condition (90° inclination), is the one calculated and used in Guidoboni et al. [41]. To assess the pressure variations associated with different tilt angles, the hydrostatic pressure component (*P*_*h*_) was computed using the previously described methodology. In the lumped 0D model, this pressure shift is represented by the hydrostatic pressure equation:

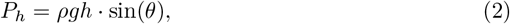

where *ρ* is the blood density, *g* is the gravitational acceleration, *h* is the vertical distance of the compartment from the reference level, and *θ* is the tilt angle. By incorporating these gravitational effects into the governing equations, the model accounts for tilt-induced changes in ocular perfusion and venous drainage, providing insights into the impact of altered gravitational conditions on retinal hemodynamics. Simulations were conducted at multiple inclination angles (0º, -6º, -15º, -30º), extending beyond the commonly used -6º HDT condition to compare with broader experimental datasets.

Changes in body position significantly impact the distribution of *P*_*h*_ throughout the cardiovascular system. This effect is particularly relevant in ocular hemodynamics, where hydrostatic shifts influence both arterial input pressure and EVP. To account for these variations, the arterial ocular pressure at a given tilt angle (*P*_in,*θ*_) is computed by adding the hydrostatic pressure component to the standing condition (90°) input pressure:

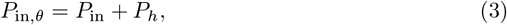

where *P*_in,*θ*_ represents the arterial input pressure at different inclination angles. This formulation ensures that the model correctly incorporates tilt-induced pressure changes affecting retinal perfusion. Furthermore, the EVP also varies with tilt angle, influencing venous outflow conditions. As described in the mathematical model by Nelson et al. [31], EVP follows the empirical relationship:

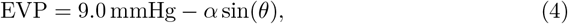

where the coefficient *α* depends on the inclination regime:

- *α* = 22.1 mmHg for *θ* ≤ 0 (e.g., HDT and inversion),
- *α* = 2.23 mmHg for *θ >* 0 (e.g., upright posture).

By incorporating these pressure-dependent boundary conditions, the model dynamically adjusts arterial and venous pressures in response to body position, improving its physiological accuracy in simulating gravitational effects on ocular hemodynamics.

For the CRA, blood velocity is governed by the following expression:

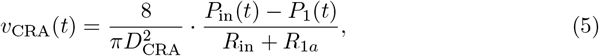

where *P*_1_(*t*) denotes the arterial downstream pressure to *P*_in_(*t*), *D*_CRA_ is the diameter of the vessel and *R*_in_ and *R*_1*a*_ represent the vascular resistances of the corresponding segment. This equation is derived from Poiseuille’s law (Table 1).

By utilizing pressure profiles corresponding to different inclination angles, key hemodynamic parameters such as peak systolic velocity (PSV), end-diastolic velocity (EDV), and mean velocity can be determined. These parameters are essential for evaluating blood flow regulation and vascular autoregulation under varying gravitational conditions, providing insights into the effects of microgravity, HDT, and other posture-induced changes on ocular circulation.

## Model Validation

The mathematical model proposed in this study consists of an improved lumped elements model defined by [31, 41]. Therefore, the model validation was performed using the reference data from each model considered. Specifically, the work of Petersen et al. The experimental campaign from the study of Petersen et al. [35] was referenced for the values of IOP and the Mean Arterial Pressure at the eye level (MAP_eye_).

As shown in Fig.2, the IOP values obtained from the experimental campaign described in Petersen et al. [35] were compared with those derived from the mathematical model presented in Nelson et al. [31]. Our simulated values (in red) successfully capture the overall trend of IOP variation with respect to the tilt angle observed in the experimental data of Petersen et al. (in black). In particular, both curves exhibit higher IOP at extreme tilt angles (i.e., -90º and 270º) and reach their minimum value around the neutral position (90º). This behavior aligns with the physiological expectation that IOP decreases when the head is roughly upright and increases when the subject is inverted or fully supine/prone.

Although the model aligns well in shape and magnitude, some local discrepancies appear. At moderate tilt angles (around ±45º), the experimental IOP deviates from the simulation, possibly reflecting individual variability in ocular physiology or measurement differences in the experimental setup. Nevertheless, our model reproduces the primary features of the IOP versus tilt relationship and provides a reliable approximation of the values reported by Petersen et al [35].

The arterial pressure at the eye level is a critical input parameter for the model, as it directly governs the hemodynamic state within the ocular system, and its averaged value is the MAP_eye_. In the present study, the baseline value at 90º was derived from an experimental measurement reported in Guidoboni et al. [41], and then an increase due to hydrostatic redistribution was applied to account for different tilt angles. This procedure allowed us to estimate the arterial pressure at the level of the eye at each inclination by considering the effect of posture on the pressure gradient between the heart and the eye.

Figure 3 shows the MAP_eye_ values computed by our model at different tilt angles (red dots) compared with the experimental data from Petersen et al. [35] (black curve with error bars). As the tilt angle deviates from the vertical (90º), arterial pressure at the eye follows a pattern consistent with hydrostatic redistribution (at extreme inclinations (e.g., -90º or 270º), it exceeds 120–130 mmHg, while near 90º, it stabilizes around 60–70 mmHg). The model-derived MAP_eye_ values show good agreement with experimental data, reinforcing the validity of the proposed approach.

**Fig 2.**
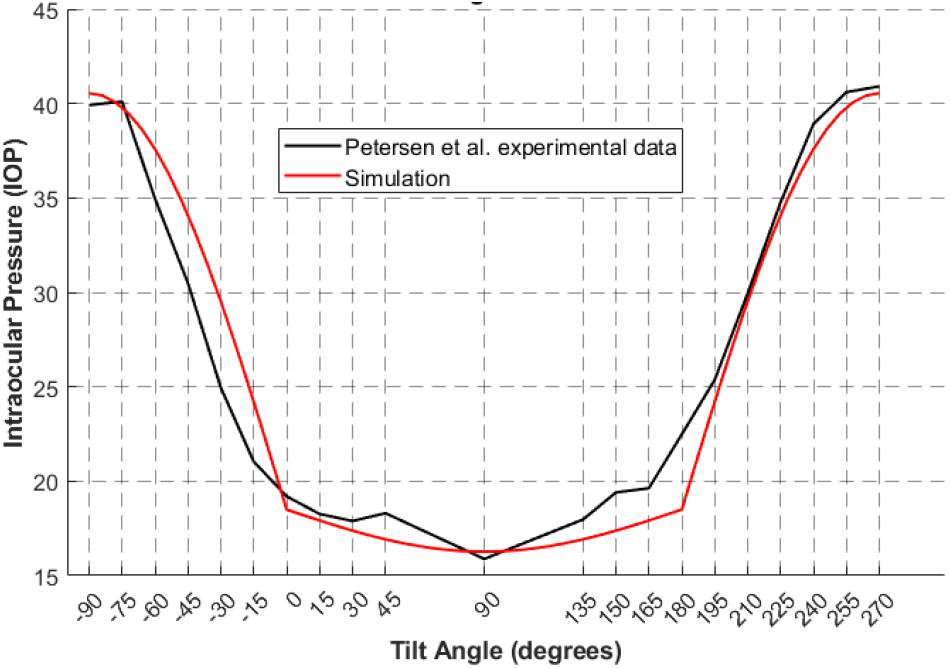
Comparison between IOP values obtained from the experimental measurements by Petersen et al. [35] (black) and those predicted by the mathematical model implemented in this study (red). The figure illustrates the IOP variation as a function of tilt angle, highlighting model accuracy and discrepancies at moderate angles.

**Fig 3.**
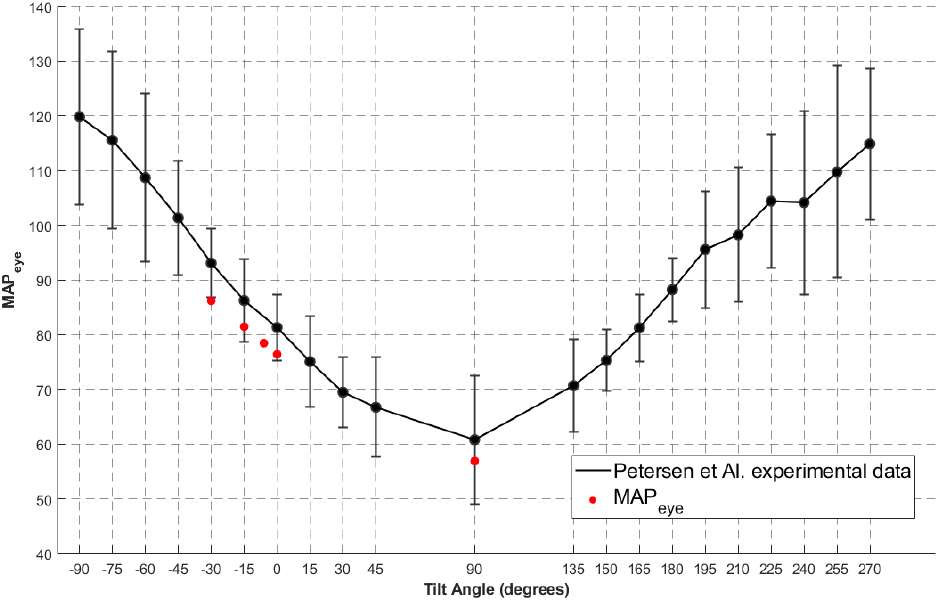
Black dashed line: Mean Arterial Pressure values from the experimental measurements from Petersen et Al. [35], with standard deviations. Scatter red points: Mean Arterial Pressure calculated from the five input pressures used in this study.

## Simulation Setup

All simulations were carried out using MATLAB^®^ (R2023a, MathWorks, Natick, MA) on a standard desktop computer. The code implements a lumped-parameter network, assembled into a system of linear equations at each time step. The governing ordinary differential equations were discretized with a fixed time step Δ*t* = 0.01 s. At each step, nodal mass-balance and compliance relations formed a linear system, which was solved using the built-in backslash operator.

## Results and discussion

### Pressures

The pressures in different retinal blood vessels were calculated for each tilt angle, showing a gradual increase in pressure as the tilt angle decreased. The highest values were observed at *θ* = −30°. The model-predicted hemodynamic changes across the CRA, arterioles, capillaries, venules, and CRV illustrate how gravitational forces influence and propagate through the retinal vasculature (Figure 4).

**Fig 4.**
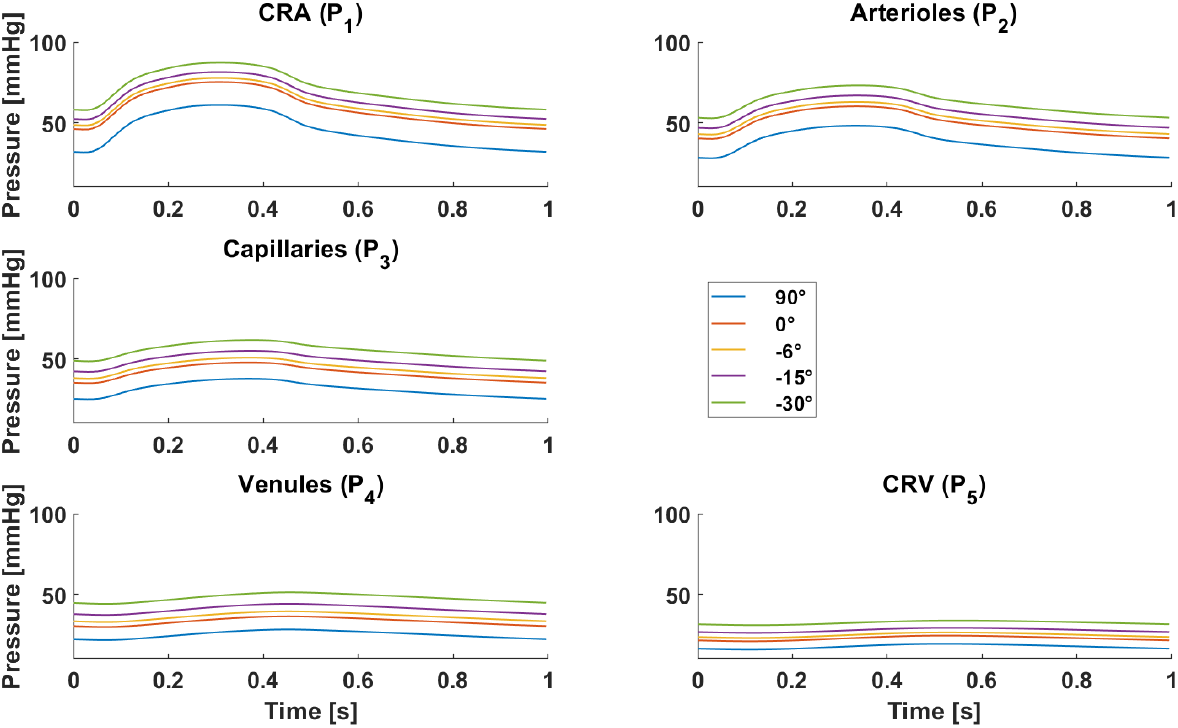
Pressure curves predicted by the mathematical model for five compartments of the retinal vasculature at different inclination angles (*θ* = 90°, 0°, −6°, −15°, −30°). A progressive increase in pressure is observed with decreasing tilt angles, with the most pronounced changes in the arterial compartments (CRA and arterioles). The legend in the figure denotes the inclination angles associated with each curve.

A distinct pattern emerges when analyzing the different vascular compartments. The arterial segments (CRA and arterioles) exhibit pronounced pulsatility, characterized by a well-defined systolic peak followed by a gradual decline, reflecting the dynamic nature of arterial blood flow regulation. As blood flow moves through the capillaries, venules, and CRV, the waveform becomes more damped, indicating the buffering effects of vascular resistance and compliance. Additionally, the baseline pressure increases as the tilt angle decreases, highlighting the role of hydrostatic pressure redistribution.

At *θ* = −30°, both peak and trough pressures are significantly higher compared to *θ* = 90°, emphasizing posture influences intraocular and retinal hemodynamics. This aligns with the expected increase in cranial hydrostatic pressure under head-down tilt conditions, which in turn affects IOP. Given the retina’s high vascularization, such alterations may compromise retinal perfusion, potentially leading to hypoperfusion, oxidative stress, and inflammation [48,49]. Understanding these mechanisms is crucial for developing countermeasures to mitigate ocular complications, including those associated with spaceflight and SANS.

These findings can be compared with the study by Guidoboni et al. [41], where simulated glaucoma conditions demonstrated a pressure increase associated with elevated IOP levels. However, unlike in that study, where IOP was directly varied, the present work shows that IOP changes as a result of body inclination.

This combined effect of increased *P*_in_(*t*) and IOP contributes to the overall rise in pressure with decreasing tilt angles. Notably, CRV pressure in the compartment reaches higher values at lower *θ*, whereas in Guidoboni et al., where *P*_in_(*t*) remained constant, CRV pressure did not deviate significantly. This suggests that the interaction between these two mechanisms plays a key role in retinal hemodynamic adaptation.

These results reinforce the importance of postural influences on ocular physiology and their potential implications for both clinical conditions and spaceflight-related adaptations.

### Velocity

The proposed model was also used to analyze the peak PSV in the CRA under different tilt angles. Figure 5 shows the numerical predictions alongside experimental data obtained from the reference study by Sirek et al. [50]. In their study, Doppler ultrasound measurements of CRA blood flow and optic nerve sheath diameter (ONSD) were conducted in both spaceflight and terrestrial conditions. Their dataset includes six non-astronaut subjects exposed to HDT at various inclinations, as well as seventeen astronauts with pre-, in-, and post-flight data from long-duration space missions aboard the ISS.

**Fig 5.**
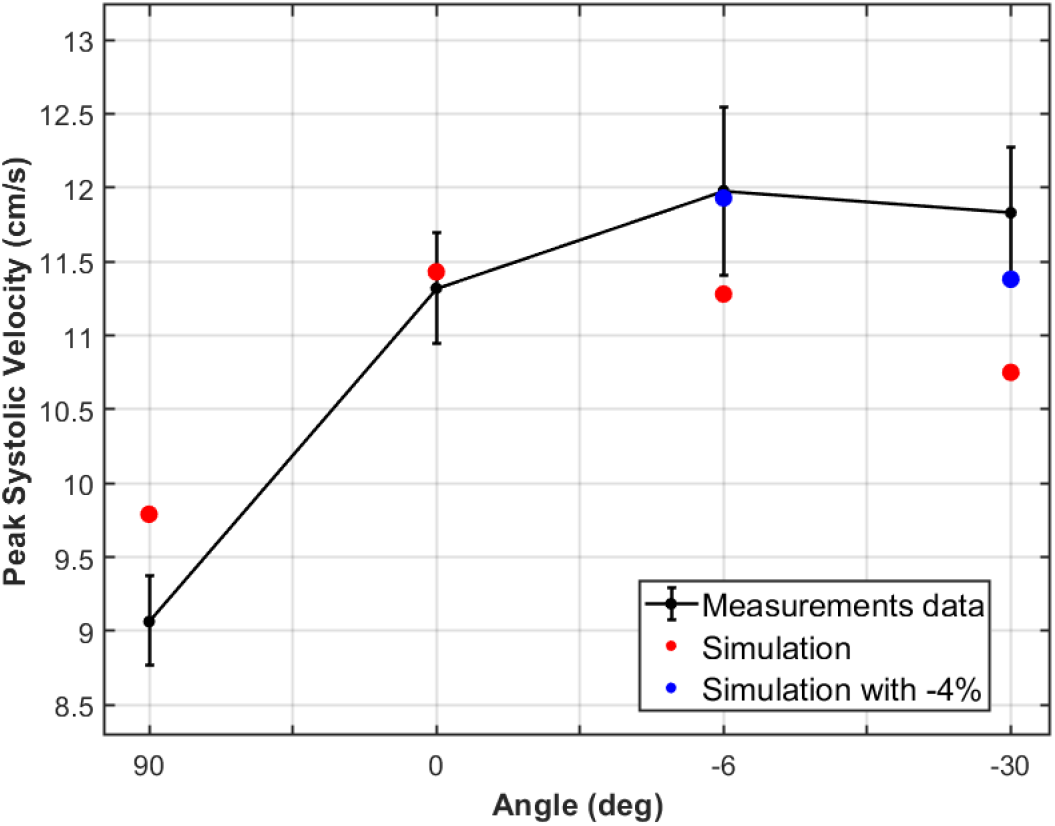
Comparison of PSV values from experimental measurements (black circles with error bars), the baseline numerical simulation (red circles), and the simulation incorporating a 4% reduction in CRA diameter (blue circles at *θ* = −6° and *θ* = −30°). The reduced-diameter condition, based on microgravity-induced vascular constriction observed by Mader et al. [13], leads to increased PSV at negative tilt angles, enhancing agreement between the numerical model and experimental data.

Additionally, a second set of simulations was performed, incorporating a 4% reduction in CRA diameter at *θ* = −6° and *θ* = −30°, motivated by the parabolic flight experiments of Mader et al. [51]. These experiments demonstrated a 4% decrease in retinal artery diameter under microgravity conditions, which was implemented in the numerical model to assess its effect on velocity predictions.

As shown in Figure 5, the numerical model closely follows the experimentally observed trend of increasing PSV with tilt angle up to *θ* = −6°, beyond which a slight decline is observed. The model’s predictions for normal CRA diameter (red markers) align well with experimental data, with deviations remaining within the range of experimental uncertainty.

For negative tilt angles (*−*6° and *−*30°), introducing the 4% diameter reduction (blue markers) results in a noticeable increase in PSV compared to the baseline simulation. This result is consistent with the hemodynamic principle that a reduction in arterial diameter increases flow velocity at a constant volumetric flow rate. Specifically, for *θ* = −6°, the PSV increased from 11.28 cm/s to 11.93 cm/s, while at *θ* = −30°, velocity increased from 10.75 cm/s to 11.38 cm/s. This modification improves agreement with experimental values, reinforcing the hypothesis that vascular constriction plays a key role in microgravity-induced hemodynamic changes.

Overall, the results demonstrate strong qualitative agreement between the numerical and experimental data. The observed sensitivity of PSV to CRA diameter varitions suggests that incorporating microgravity-induced vascular responses is essential for accurately simulating retinal hemodynamics.

### Retinal Blood Flow

By applying the governing equation at different tilt angles (*θ* = 90°, 0°, −6°, −30°), we computed the retinal blood flow and related it to the corresponding OPP, introduced in Table 1.

Garhofer et al. [52] measured total retinal blood flow in 64 healthy volunteers and found no significant correlation between retinal blood flow and MAP, IOP, or OPP, suggesting that autoregulation stabilizes flow despite blood pressure variations.

In Fig. 6, we compare our simulated retinal arterial flow values (red: no reduction, blue: 4% reduction) with the experimental data from Garhofer et al. [52], which correspond to healthy subjects in the upright position (*θ* = 90°). Our model aligns well with the observed variability, particularly at lower OPP values. As OPP increases (corresponding to tilt angles of 0°, −6°, and −30°, where MAP is generally higher), the predicted flow remains within the experimental range.

**Fig 6.**
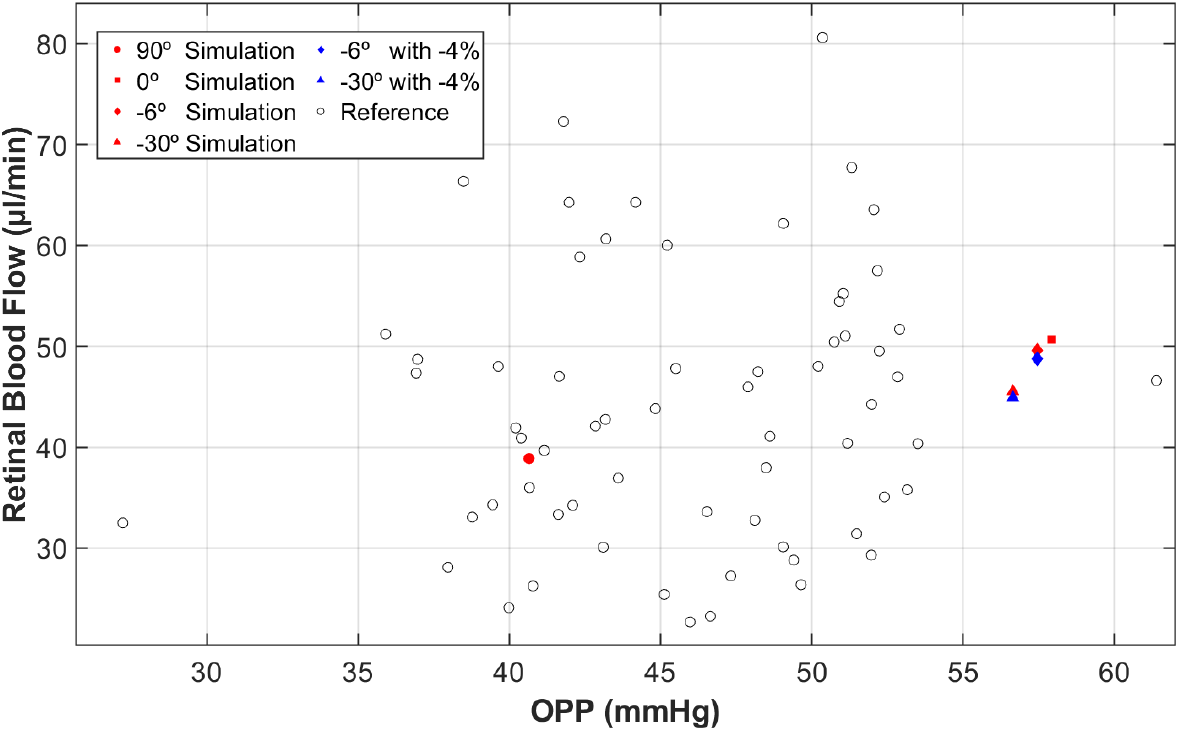
Retinal blood flow as a function of OPP, comparing simulation results (red and blue dots) with experimental data from Garhofer et al. [52].

These results reinforce the hypothesis that retinal circulation is effectively autoregulated across a range of inclinations and perfusion pressures. Despite variations in MAP due to postural changes, the simulated blood flow remains relatively stable, consistent with the experimental data reported by Garhofer et al. [52].

Figure 7 also presents trends in retinal blood flow, PSV, EDV, and the RI as functions of *θ*. As *θ* increases (moving from −30° toward 90°), retinal blood flow progressively rises, in parallel with an increase in both PSV and EDV. Specifically, PSV increases from approximately 8 cm/s at *θ* = −30° to nearly 10 cm/s at *θ* = 90°, while EDV nearly doubles its initial value.

**Fig 7.**
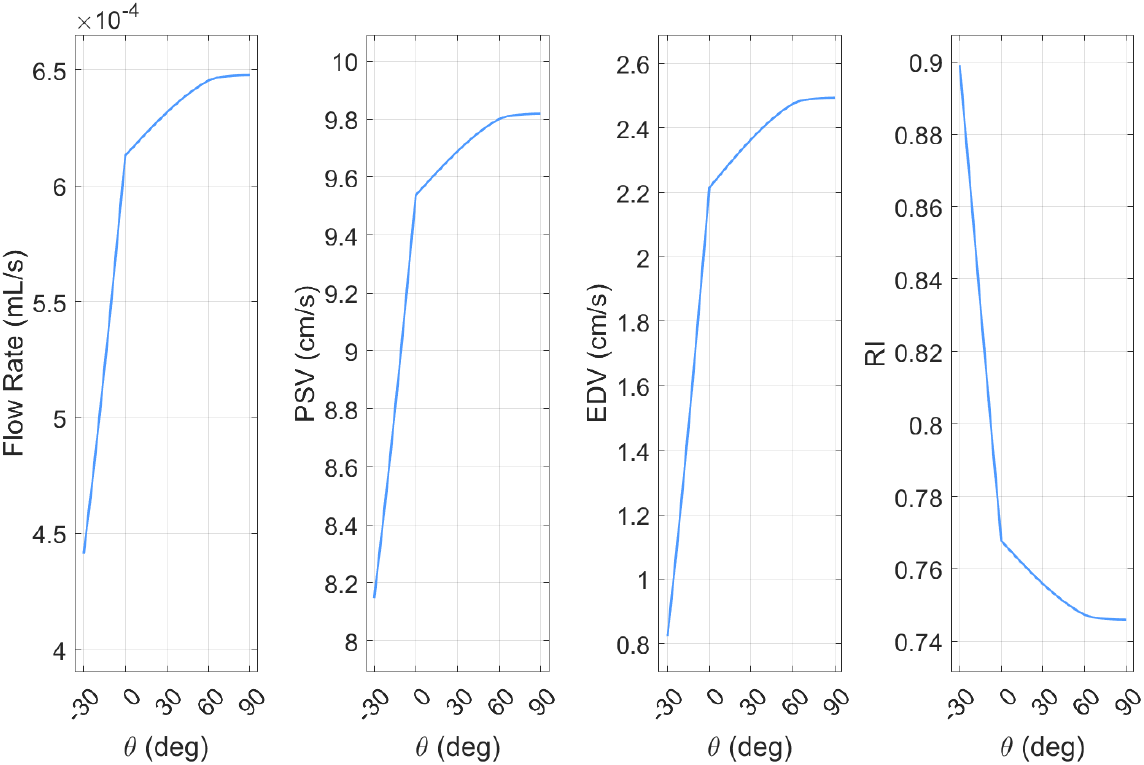
Variation of retinal hemodynamic parameters as a function of tilt angle (*θ*). (Left to right): Retinal blood flow rate, PSV, EDV, and RI.

This trend affects RI, which decreases as *θ* increases. The rise in EDV significantly reduces the difference (PSV − EDV), leading to a lower RI. Thus, tilting from an upright position to −30° results in reduced retinal blood flow, lower PSV and EDV, and an increased vascular resistance index. These changes likely reflect the influence of the hydrostatic gradient and local hemodynamic adjustments.

## Limitations and future developments

While the current model provides valuable insights into retinal hemodynamics under altered gravity, it has several limitations. A key constraint is its inability to fully replicate spaceflight conditions, as factors like chronic fluid redistribution, cardiovascular adaptations, and prolonged microgravity exposure are not explicitly modeled [2,19,24,53]. Although HDT serves as a terrestrial analog, it does not fully capture the sustained interaction between ICP and retinal circulation. Additionally, the model lacks individual physiological variability, including biological sex differences, pre-existing conditions, and personalized hemodynamic responses, which could significantly impact ocular adaptation [54–56]. Another limitation is the exclusion of radiation-induced retinal effects, a crucial aspect of space medicine that may interact with vascular dynamics in ways not yet explored [57–59].

To enhance physiological relevance and predictive capability, future developments will integrate the retinal circulation framework with established models of cerebral hemodynamics and CSF dynamics [30, 60, 61]. A more detailed venous system will be incorporated to better understand thrombosis formation, blood distribution, and vascular compliance in response to gravitational changes [16, 62–65]. Expanding the model to clinical conditions such as hypertensive retinopathy and glaucoma will further test its versatility and clinical impact [41]. Lastly, integrating autoregulatory mechanisms and optimization techniques will improve patient-specific parameter estimation, enhancing predictive accuracy [27, 66, 67]. These advancements will strengthen the role of the model in studying both SANS and terrestrial pathologies associated with altered intracranial and intraocular pressures.

## Conclusion

This study introduces an innovative computational model for analyzing retinal hemodynamic responses under varying gravitational and postural conditions. By incorporating dynamic IOP and gravitational effects, the computational model developed provides a physiologically accurate representation of ocular circulation, addressing limitations of previous approaches that treated IOP as a fixed parameter. Validation against experimental data has demonstrated the model’s ability to accurately reproduce changes in ocular pressure and retinal blood flow observed under tilt and microgravity conditions, reinforcing its applicability in studying SANS.

Despite its advancements, the model presents some challenges concerning to the capacity to simulate prolonged microgravity exposure, individual physiological variability, and radiation-induced ocular changes. Future research should focus on integrating this framework with cerebral hemodynamic and CFS dynamics, improving the representation of vascular autoregulatory mechanisms, and personalizing the model parameters based on clinical data. These improvements will not only enhance predictions of ocular alterations in astronauts but also extend the model’s applicability to terrestrial pathologies related to intracranial and intraocular pressure dysfunctions, such as glaucoma and hypertensive retinopathy. Ultimately, this model represents a significant step forward in simulating ocular physiology in altered environments, laying the foundation for the development of effective countermeasures against spaceflight-related visual risks and providing a valuable tool for research in ophthalmology and applied neuroscience.

## Appendix

### Mathematical model

The numerical solution of the differential equations governing *P*_1_, *P*_2_, *P*_4_, *P*_5_, and IOP was implemented using a forward Euler time discretization scheme combined with a matrix-based solution for the linearized equations at each time step. Specifically, the method constructs a linear system *A* · *P*_new_ = *B* for the state variables at each time step, where *A* is a coefficient matrix dependent on system parameters (e.g., resistances, compliances), and *B* incorporates terms from the previous time step. The backslash operator (\) in MATLAB is employed to efficiently solve this system.

The dynamic behavior of intraocular pressure and retinal circulatory pressures (CRA, arterioles, venules, CRV) is modeled in conjunction with vascular compliance and resistances. The resulting time series provide a detailed temporal profile of the pressure dynamics, capturing physiological phenomena such as pulsatility and flow regulation in response to systemic and episcleral venous pressures. The vascular resistances (*R*) and capacitances (*C*) introduced by Guidoboni er al. [41] are summarized and reported in Table 3 . These parameters play a crucial role in modeling the retinal vasculature, capturing the interplay between pressure, flow, and vessel elasticity. While some resistances are constant, others are variable, dynamically adapting to changes in transmural pressure differences (Δ*P*_*t*_), flow, or autoregulatory mechanisms. The reported values represent baseline or steady-state conditions under normal physiological parameters. In Table 4 the key physiological parameters included in the model, introduced by Nelson et al. [31] regarding the the IOP dynamics, are reported.

**Table 2.** Summary of simulated hemodynamic parameters across various tilt angles. Maximum and minimum pressures in the retinal compartments (P1-P5), IOP, and OPP demonstrate consistent changes with decreasing tilt angles, indicating significant gravitational influence on ocular hemodynamics

|  | 90° | 0° | -6° | -15° | -30° |
| --- | --- | --- | --- | --- | --- |
| MAPeye (mmHg) | 56.92 | 76.42 | 78.46 | 81.46 | 86.17 |
| P1max - P1min (mmHg) | 61.08 - 31.47 | 75.30 - 45.67 | 77.82 - 48.17 | 81.55 - 51.85 | 87.34 - 57.60 |
| P2max - P2min (mmHg) | 48.30 - 27.86 | 60.25 - 39.79 | 62.99 - 42.47 | 67.01 - 46.43 | 73.25 - 52.60 |
| P3max - P3min (mmHg) | 37.77 - 24.75 | 47.83 - 34.77 | 50.75 - 37.61 | 55.05 - 41.81 | 61.73 - 48.35 |
| P4max - P4min (mmHg) | 28.03 - 21.47 | 36.18 - 29.57 | 39.32 - 32.57 | 43.93 - 36.99 | 51.07 - 43.88 |
| P5max - P5min (mmHg) | 19.08 - 15.59 | 24.24 - 20.76 | 26.17 - 22.85 | 29.04 - 25.93 | 33.54 - 30.75 |
| IOP (mmHg) | 16.27 | 18.50 | 20.80 | 24.20 | 29.52 |
| OPP (mmHg) | 40.65 | 57.92 | 57.66 | 57.46 | 56.65 |

**Table 3.** Vascular resistances and compliances used in the numerical model, adapted from Guidoboni et al. [41].

| Parameter | Value | Units |
| --- | --- | --- |
| **Resistances ($R$)** |  |  |
| Input resistance to CRA | $2.25 \times 10^4$ | mmHg·s/mL |
| CRA retrobulbar segment | $4.30 \times 10^3$ | mmHg·s/mL |
| CRA translaminar segment | $1.96 \times 10^2$ | mmHg·s/mL |
| CRA intraocular segment | $9.78 \times 10^2$ | mmHg·s/mL |
| Arterioles | $6.00 \times 10^3$ | mmHg·s/mL |
| Capillaries | $5.68 \times 10^3$ | mmHg·s/mL |
| Venules | $3.11 \times 10^3$ | mmHg·s/mL |
| CRV intraocular segment | $1.35 \times 10^3$ | mmHg·s/mL |
| CRV retrobulbar segment | $6.15 \times 10^3$ | mmHg·s/mL |
| Output resistance | $5.74 \times 10^3$ | mmHg·s/mL |
| **Capacitances ($C$)** |  |  |
| CRA compliance ($C_1$) | $7.22 \times 10^{-7}$ | mL/mmHg |
| Arteriolar compliance ($C_2$) | $7.53 \times 10^{-7}$ | mL/mmHg |
| Venular compliance ($C_4$) | $1.67 \times 10^{-5}$ | mL/mmHg |
| CRV compliance ($C_5$) | $1.07 \times 10^{-5}$ | mL/mmHg |

**Table 4.** Physiological parameters involved in the IOP dynamics, from Nelson et al. [31].

| Parameter | Value | Units |
| --- | --- | --- |
| **Aqueous Humor Dynamics** |  |  |
| Rate of aqueous humor inflow ($Q_{aq,in}$) | $2.4 \times 10^{-3}$ | mL/min |
| Trabecular outflow facility ($C_{tm}$) | $0.30 \times 10^{-3}$ | mL/min/mmHg |
| Uveoscleral outflow rate ($Q_{uv}$) | $0.4 \times 10^{-3}$ | mL/min |
| **Compliance Parameters** |  |  |
| Compliance of retrobulbar space ($C_{rg}$) | $1.1 \times 10^{-6}$ | mL/mmHg |
| **Other Variables** |  |  |
| Volume of globe ($V_g$) | 6.5 | mL |
| Cerebrospinal fluid pressure ($P_{csf}$) | 13 | mmHg |

#### Pressure Dynamics Equations

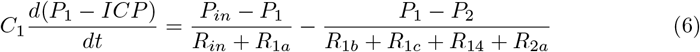

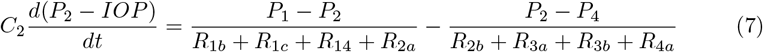

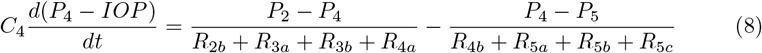

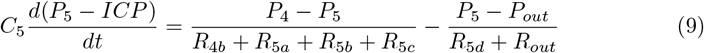

#### Intraocular Pressure (IOP) Equation

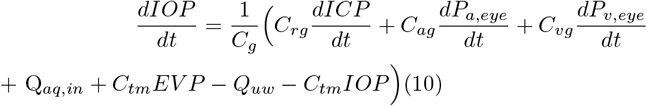

#### rSAS-to-Globe Compliance (*Crg*)

Compliance between the retrobulbar subarachnoid space (rSAS) and the globe, representing the influence of cerebrospinal fluid pressure (Pcsf) on IOP.

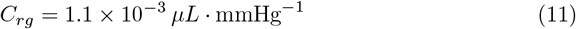

#### Total Globe Compliance (*Cg*)

Total globe compliance represents the overall compliance of the eye, including the corneoscleral shell, rSAS, and blood-to-globe compliance.

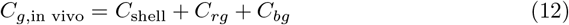

where *C*_*bg*_ = *C*_*ag*_ + *C*_*vg*_ is the net blood-to-globe compliance.

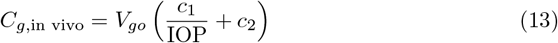

with empirical constants:

- *c*_1_ = 4.87 *×* 10^*−*3^
- *c*_2_ = 3.90 *×* 10^*−*5^ mmHg^*−*1^
- *V*_*go*_ is the initial globe volume of 6, 500 *µ*L

#### Blood-to-Globe Compliance (*C*_*bg*_)

Blood-to-globe compliance (*C*_*bg*_) represents the compliance of the ocular blood vessels contributing to intraocular pressure dynamics. It is modeled similarly to the total globe compliance using empirical constants *c*_1_ and *c*_2_:

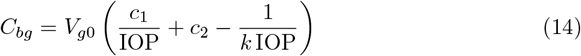

where *k*, the nondimensional globe stiffness, is calculated by multiplying Friedenwald’s ocular rigidity coefficient *K* = 0.048 *µL*^*−*1^ by the initial globe volume *V*_*g*0_, resulting in *k* = 312.

#### Arterial Blood-to-Globe Compliance (*Cag*)

Compliance of arterial blood vessels contributing to the overall blood-to-globe compliance.

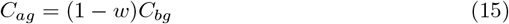

where *w* = 0.7 is the fraction of blood distal to the arteries, approximately at venous pressure.

#### Venous Blood-to-Globe Compliance (*Cvg*)

Compliance of venous blood vessels contributing to the overall blood-to-globe compliance.

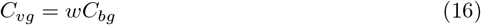

#### Trabecular Meshwork Compliance (*Ctm*)

Aqueous outflow facility through the trabecular meshwork, influencing intraocular pressure.

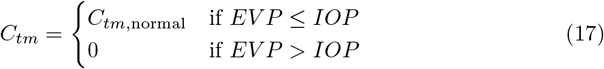

where:

- *C*_*tm*,normal_ = 0.30 *µL ·* min^*−*1^ *·* mmHg^*−*1^
- *EV P* is the episcleral venous pressure.

#### Aqueous Humor Dynamics

Aqueous humor inflow (*Q*_aq,in_) and outflow (*Q*_aq,out_) are modeled as:

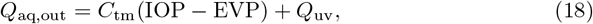

where *C*_tm_ is the trabecular meshwork outflow facility, EVP the episcleral venous pressure, and *Q*_uv_ the uveoscleral flow rate.

#### Venous Pressure Regulation and Tilt-Induced Variations

The relationship between EVP and CVP, which governs venous outflow and is influenced by body tilt.

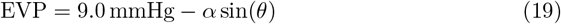

where *α* is an empirically determined coefficient that varies with inclination angle:

This equation models the dependence of EVP on body tilt, reflecting gravitational effects on venous drainage. Under HDT, EVP increases due to elevated venous hydrostatic pressure, while in the upright position, EVP remains lower.

##### Venous Outflow Regulation

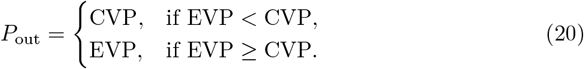

This formulation ensures that venous outflow remains physiologically viable under different gravitational conditions. When EVP exceeds CVP, venous pressure follows EVP; otherwise, it remains equal to CVP. This regulatory mechanism prevents venous collapse and maintains adequate ocular circulation, particularly under altered gravitational environments such as microgravity or HDT.

##### Arterial Input Pressure and Hydrostatic Effects

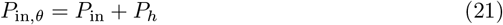

where

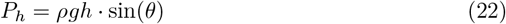

accounts for the hydrostatic pressure shift due to body inclination. This adjustment ensures that arterial input pressure accurately reflects gravitational effects, improving the physiological realism of the model in simulating ocular hemodynamics across different tilt conditions.

#### General Resistance Model

Variable resistances (*R*) are calculated using Poiseuille’s law, modified to account for the vessel’s deformation and flow dynamics:

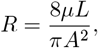

where:

- *µ*: fluid dynamic viscosity,
- *L*: vessel length,
- *A*: cross-sectional area of the vessel.

Changes in *A* are governed by vessel wall properties and external pressures.

#### Passive Variable Resistances

Passive resistances are influenced by the transmural pressure difference (Δ*P*_*t*_ = *P*_internal_ *− P*_external_) and vessel compliance. For veins and intraocular vessels:

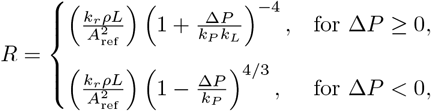

*k*_*P*_ , *k*_*L*_, **and** *k*_*R*_

These parameters are essential for modeling the mechanical properties of vessels, particularly their compliance and resistance to pressure changes:

- *k*_*P*_ **: Pressure Stiffness Coefficient** Represents the stiffness of the vessel wall under positive or negative transmural pressure (Δ*P*_*t*_). Defined as:

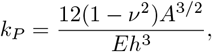

where:
  - *E*: Young’s modulus of the vessel wall, reflecting its elasticity,
  - *h*: vessel wall thickness,
  - *ν*: Poisson’s ratio, indicating the degree of material compressibility,
  - *A*: cross-sectional area of the vessel.
- *k*_*L*_**: Length Stiffness Coefficient** Governs the resistance to flow through the vessel by incorporating the geometrical effects of length and deformation. Defined as:

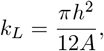

where:

- *A*: cross-sectional area of the vessel,
- *h*: vessel wall thickness.
- *k*_*R*_**: Resistance Scaling Factor** Accounts for fluid viscosity (*µ*) and the geometric properties of the vessel, affecting the baseline resistance. Defined as:

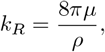

where:

- *µ*: dynamic viscosity of blood,
- *ρ*: blood density.
- **Resistances (***R*):
- *R*_in_ represents the resistance between the systemic arterial supply and the central retinal artery (CRA).
  - *R*_1*a*_, *R*_1*b*_, *R*_1*c*_, and *R*_1*d*_ correspond to the resistances in the retrobulbar, translaminar, and intraocular segments of the CRA. Among these, *R*_1*c*_ and *R*_1*d*_ are variable, adjusting dynamically to changes in pressure.
  - *R*_2*a*_, *R*_2*b*_ describe autoregulated resistances in the arterioles, reflecting their capacity to adapt to local metabolic demands.
  - *R*_3*a*_, *R*_3*b*_ represent constant resistances in the retinal capillaries.
  - *R*_4*a*_, *R*_4*b*_ capture variable resistances in the venular segments.
  - *R*_5*a*_, *R*_5*b*_, *R*_5*c*_, and *R*_5*d*_ refer to resistances in the intraocular and retrobulbar segments of the central retinal vein (CRV). The retrobulbar resistance (*R*_5*d*_) is constant, whereas the others are variable.
  - *R*_out_ represents the resistance between the CRV and systemic venous drainage.
- **Compliances (***C*):
  - *C*_1_ denotes the compliance of the CRA.
  - *C*_2_ captures the compliance of the arterioles.
  - *C*_4_ and *C*_5_ reflect the compliance of the venules and the CRV, respectively.

## Acknowledgements

The authors acknowledge financial support from the Spanish Ministry of Economy and Competitiveness, through the support of “Proyecto CPP2021-008546 financiado por MCIN/AEI/10.13039/501100011033” por la Unión Europea NextGeneration EU/PRTR” and “Proyecto PID2021-122518OB-I00 financiado por MCIN/AEI/10.13039/501100011033” por FEDER Una manera de hacer Europa”.

## Author contributions

**Conceptualization**: Michele Nigro, Eduardo Soudah. **Methodology**: Michele Nigro, Eduardo Soudah. **Formal Analysis**: Michele Nigro. **Software**:Michele Nigro, Andrea Montanino. **Validation**: Michele Nigro, Andrea Montanino. **Methodology**: Michele Nigro, Eduardo Soudah. **Funding acquisition**: Eduardo Soudah. **Writing – original draft**:Michele Nigro, Andrea Montanino, Eduardo Soudah. **Writing – review editing**: Michele Nigro, Andrea Montanino, Eduardo Soudah.

## References

1. Aeronautics N, (NASA) SA. Human Research Roadmap; 2020. Available from: https://humanresearchroadmap.nasa.gov.

2. Galdamez LA, Mader TH, Ong J, Kadipasaoglu CM, Lee AG. A multifactorial, evidence-based analysis of pathophysiology in Spaceflight Associated Neuro-Ocular Syndrome (SANS). Eye. 2025; p. 1–10.

3. Mader THea. Optic Disc Edema, Globe Flattening, and Choroidal Folds after Long-Duration Space Flight. Ophthalmology. 2011;118:2058–2069. doi:10.1016/j.ophtha.2011.06.021.

4. Lee AG, Mader TH, Gibson CR, Tarver W, Rabiei P, Riascos RF, et al. Space-flight associated neuro-ocular syndrome (SANS) and the neuro-ophthalmologic effects of microgravity: a review and an update. npj Microgravity. 2020;6:7. doi:10.1038/s41526-020-0097-9.

5. Nelson ES, Mulugeta L, Myers JG. Microgravity-induced fluid shift and ophthalmic changes. Life. 2014;4(4):621–665.

6. Taibbi G, Cromwell RL, Kapoor KG, Godley BF, Vizzeri G. The Effect of Microgravity on Ocular Structures and Visual Function: A Review. Survey of Ophthalmology. 2013;58(2):155–163. doi:10.1016/j.survophthal.2012.04.002.

7. Killer H, Jaggi G, Flammer J, Miller N, Huber A. The optic nerve: a new window into cerebrospinal fluid composition? Brain. 2006;129(4):1027–1030.

8. Wostyn P, Mader TH, Gibson CR, Killer HE. The perivascular space of the central retinal artery as a potential major cerebrospinal fluid inflow route: implications for optic disc edema in astronauts. Eye. 2020;34(4):779–780.

9. Zhang LF, Hargens AR. Spaceflight-induced intracranial hypertension and visual impairment: pathophysiology and countermeasures. Physiological reviews. 2018;98(1):59–87.

10. Barisano G, Sepehrband F, Collins HR, Jillings S, Jeurissen B, Taylor JA, et al. The effect of prolonged spaceflight on cerebrospinal fluid and perivascular spaces of astronauts and cosmonauts. Proceedings of the National Academy of Sciences (PNAS). 2022;119(17):e2120439119. doi:10.1073/pnas.2120439119.

11. Laurie SS, Macias BR, Dunn JT, Young M, Stern C, Lee SM, et al. Optic disc edema after 30 days of strict head-down tilt bed rest. Ophthalmology. 2019;126(3):467–468.

12. Mader TH, Gibson CR, Otto CA, Sargsyan AE, Miller NR, Subramanian PS, et al. Persistent asymmetric optic disc swelling after long-duration space flight: implications for pathogenesis. Journal of Neuro-Ophthalmology. 2017;37(2):133– 139.

13. Mader TH, Gibson CR, Pass AF, Kramer LA, Lee AG, Fogarty J, et al. Optic disc edema, globe flattening, choroidal folds, and hyperopic shifts observed in astronauts after long-duration space flight. Ophthalmology. 2011;118(10):2058– 2069.

14. Ong J, Tarver W, Brunstetter T, Mader TH, Gibson CR, Mason SS, et al. Spaceflight associated neuro-ocular syndrome: proposed pathogenesis, terrestrial analogues, and emerging countermeasures. British Journal of Ophthalmology. 2023;107(7):895–900.

15. Lee AG, Tarver WJ, Mader TH, Gibson CR, Hart SF, Otto CA. Neuro-ophthalmology of space flight. Journal of Neuro-Ophthalmology. 2016;36(1):85–91.

16. Whittle RS, Diaz-Artiles A. Gravitational Effects on Carotid and Jugular Characteristics in Graded Head-Up and Head-Down Tilt. Journal of Applied Physiology. 2023;134:217–229. doi:10.1152/japplphysiol.00248.2022.

17. Lerner DJ, Chima RS, Patel K, Parmet AJ. Ultrasound guided lumbar puncture and remote guidance for potential in-flight evaluation of VIIP/SANS. Aerospace Medicine and Human Performance. 2019;90(1):58–62.

18. Hiles LA, Donoviel DB, Bershad EM. Noninvasive brain physiology monitoring for extreme environments: a critical review. Journal of Neurosurgical Anesthesiology. 2015;27(4):318–328.

19. Watenpaugh DE. Analogs of microgravity: head-down tilt and water immersion. Journal of Applied Physiology. 2016;120(8):904–914.

20. Ong J, Lee AG, Moss HE. Head-down tilt bed rest studies as a terrestrial analog for spaceflight associated neuro-ocular syndrome. Frontiers in Neurology. 2021;12:648958.

21. Linder BJ, Trick GL, Wolf ML. Altering body position affects intraocular pressure and visual function. Investigative Ophthalmology Visual Science. 1988;29(10):1492– 1497.

22. Hargens AR, Vico L. Long-duration bed rest as an analog to microgravity. Journal of Applied Physiology. 2016;120:891–903. doi:10.1152/japplphysiol.00935.2015.

23. Van Akin MP, Lantz OM, Fellows AM, Toutain-Kidd C, Zegans M, Buckey JC, et al. Acute effects of postural changes and lower body positive and negative pressure on the eye. Frontiers in Physiology. 2022;13:933450. doi:10.3389/fphys.2022.933450.

24. Barkaszi I, Ehmann B, Tölgyesi B, Balázs L, Altbäcker A. Are head-down tilt bedrest studies capturing the true nature of spaceflight-induced cognitive changes? A review. Frontiers in Physiology. 2022;13:1008508. doi:10.3389/fphys.2022.1008508.

25. Caddy HT, Kelsey LJ, Parker LP, Green DJ, Doyle BJ. Modelling large scale artery haemodynamics from the heart to the eye in response to simulated microgravity. npj Microgravity. 2024;10:7. doi:10.1038/s41526-024-00348-w.

26. Heldt T, Shim EB, Kamm RD, Mark RG. Computational modeling of cardiovascular response to orthostatic stress. Journal of Applied Physiology. 2002;92:1239–1254. doi:10.1152/japplphysiol.00241.2001.

27. Heldt T. Computational models of cardiovascular response to orthostatic stress. Massachusetts Institute of Technology; 2004.

28. Fois M, Maule SV, Giudici M, Valente M, Ridolfi L, Scarsoglio S. Cardiovascular response to posture changes: multiscale modeling and in vivo validation during head-up tilt. Frontiers in Physiology. 2022;13:826989.

29. Nelson ES, Myers Jr JG, Lewandowski BE, Ethier CR, Samuels BC. Acute effects of posture on intraocular pressure. PLoS One. 2020;15(2):e0226915.

30. Scarsoglio S, Fois M, Ridolfi L. Increased Hemodynamic Pulsatility in the Cerebral Microcirculation During Parabolic Flight-Induced Microgravity: A Computational Investigation. Acta Astronautica. 2023;211:344–352. doi:10.1016/j.actaastro.2023.06.018.

31. Nelson ES, Mulugeta L, Feola A, Raykin J, Myers JG, Samuels BC, et al. The impact of ocular hemodynamics and intracranial pressure on intraocular pressure during acute gravitational changes. Journal of Applied Physiology. 2017;123:352– 363. doi:10.1152/japplphysiol.00102.2017.

32. Salerni F, Repetto R, Harris A, Pinsky P, Prud’homme C, Szopos M, et al. Biofluid modeling of the coupled eye-brain system and insights into simulated microgravity conditions. PLoS ONE. 2019;14(8):e0216012. doi:10.1371/journal.pone.0216012.

33. Fois M, Diaz-Artiles A, Zaman SY, Ridolfi L, Scarsoglio S. Linking cerebral hemodynamics and ocular microgravity-induced alterations through an in silico-in vivo head-down tilt framework. npj Microgravity. 2024;10(22). doi:10.1038/s41526024-00366-8.

34. Wilson SL, Schulte KM, Steins A, Gruen RL, Tucker EM, van Loon LM. Computational modeling of heart failure in microgravity transitions. Frontiers in Physiology. 2024;15:1351985.

35. Petersen LG, Whittle RS, Lee JH, Sieker J, Carlson J, Finke C, et al. Gravitational effects on intraocular pressure and ocular perfusion pressure. Journal of Applied Physiology. 2022;132:24–35. doi:10.1152/japplphysiol.00546.2021.

36. Navasiolava N, Yuan M, Murphy R, Robin A, Coupé M, Wang L, et al. Vascular and microvascular dysfunction induced by microgravity and its analogs in humans: mechanisms and countermeasures. Frontiers in physiology. 2020;11:952.

37. Grigoryan EN. Impact of Microgravity and Other Spaceflight Factors on Retina of Vertebrates and Humans In Vivo and In Vitro. Life. 2023;13:1263. doi:10.3390/life13061263.

38. Iftime A, Tofolean IT, Pintilie V, Călinescu O, Busnatu S, Papacocea IR. Differential Functional Changes in Visual Performance during Acute Exposure to Microgravity Analogue and Their Potential Links with Spaceflight-Associated Neuro-Ocular Syndrome. Diagnostics. 2024;14(17):1918.

39. Masland RH. The fundamental plan of the retina. Nature neuroscience. 2001;4(9):877–886.

40. Baden T. The vertebrate retina: a window into the evolution of computation in the brain. Current Opinion in Behavioral Sciences. 2024;57:101391.

41. Guidoboni G, Harris A, Cassani S, Arciero J, Siesky B, Amireskandari A, et al. Intraocular Pressure, Blood Pressure, and Retinal Blood Flow Autoregulation: A Mathematical Model to Clarify Their Relationship and Clinical Relevance. Investigative Ophthalmology Visual Science. 2014;55:4105–4118. doi:10.1167/iovs.13-13611.

42. Westerhof N, Bosman F, De Vries CJ, Noordergraaf A. Analog Studies of the Human Systemic Arterial Tree. Biomedwarm. 1969;2:121–143.

43. Garber L, Khodaei S, Reymond P, Stergiopulos N. The Critical Role of Lumped Parameter Models in Patient-Specific Hemodynamics. Journal of Biomechanical Engineering. 2023;145(8):081004.

44. Diaz-Artiles A, Heldt T, Young LR. Computational model of cardiovascular response to centrifugation and lower body cycling exercise. Journal of Applied Physiology. 2019;127:1453–1468. doi:10.1152/japplphysiol.00314.2019.

45. Ghitti B, Toro EF, Müller LO. Nonlinear Lumped-Parameter Models for Blood Flow Simulations in Networks of Vessels. ArXiv. 2022;.

46. Brown AG, Shi Y, Marzo A, Staicu C, Valverde I, Beerbaum P, et al. Accuracy vs. computational time: Translating aortic simulations to the clinic. Journal of Biomechanics. 2012;45:516–523. doi:10.1016/j.jbiomech.2011.11.041.

47. Kokalari I, Karaja T, Guerrisi M. Review on lumped parameter method for modeling the blood flow in systemic arteries. Journal of Biomedical Science and Engineering. 2013;6:92–99. doi:10.4236/jbise.2013.61012.

48. Di Marco E, Aiello F, Lombardo M, Di Marino M, Missiroli F, Mancino R, et al. A literature review of hypertensive retinopathy: systemic correlations and new technologies. European Review for Medical and Pharmacological Sciences. 2022;26(18):6424–6443.

49. Modi P, Arsiwalla T. Hypertensive retinopathy. In: StatPearls [Internet]. StatPearls Publishing; 2023.

50. Sirek AS, Garcia K, Foy M, Ebert D, Sargsyan A, Wu JH, et al. Doppler ultrasound of the central retinal artery in microgravity. Aviation, Space, and Environmental Medicine. 2014;85:3–8. doi:10.3357/ASEM.3750.2014.

51. Mader TH, Gibson CR, Caputo M, Hunter N, Taylor G, Charles J, et al. Intraocular Pressure and Retinal Vascular Changes During Transient Exposure to Microgravity. American Journal of Ophthalmology. 1993;115:347–350.

52. Garhofer G, Werkmeister R, Dragostinoff N, Schmetterer L. Retinal Blood Flow in Healthy Young Subjects. Investigative Ophthalmology Visual Science. 2012;53:698– 703. doi:10.1167/iovs.11-8624.

53. Oluwafemi FA, Neduncheran A. Analog and simulated microgravity platforms for life sciences research: Their individual capacities, benefits and limitations. Advances in Space Research. 2022;69(7):2921–2929.

54. Kordi M, Kluge N, Kloeckner M, Russomano T. Gender influence on the performance of chest compressions in simulated hypogravity and microgravity. Aviation, Space, and Environmental Medicine. 2012;83(7):643–648.

55. Evans JM, Knapp CF, Goswami N. Artificial gravity as a countermeasure to the cardiovascular deconditioning of spaceflight: gender perspectives. Frontiers in physiology. 2018;9:716.

56. Strock N, Rivas E, Goebel KM. The effects of space flight and microgravity exposure on female astronaut health and performance. In: 2023 IEEE Aerospace Conference. IEEE; 2023. p. 01–12.

57. Mao XW, Stanbouly S, Chieu B, Sridharan V, Allen AR, Boerma M. Low dose space radiation-induced effects on the mouse retina and blood-retinal barrier integrity. Acta Astronautica. 2022;199:412–419.

58. Waisberg E, Ong J, Paladugu P, Kamran SA, Zaman N, Tavakkoli A, et al. Radiation-induced ophthalmic risks of long duration spaceflight: Current investigations and interventions. European Journal of Ophthalmology. 2024;34(5):1337– 1345.

59. Cialdai F, Bolognini D, Vignali L, Iannotti N, Cacchione S, Magi A, et al. Effect of space flight on the behavior of human retinal pigment epithelial ARPE-19 cells and evaluation of coenzyme Q10 treatment. Cellular and Molecular Life Sciences. 2021;78:7795–7812.

60. Holmlund P, Støverud KH, Eklund A. Mathematical Modelling of the CSF System: Effects of Microstructures and Posture on Optic Nerve Subarachnoid Space Dynamics. Fluids and Barriers of the CNS. 2022;19:67. doi:10.1186/s12987-022-00366-4.

61. Ursino M, Giannessi M. A model of cerebrovascular reactivity including the circle of Willis and cortical anastomoses. Annals of biomedical engineering. 2010;38:955– 974.

62. Yu DY, Cringle SJ, Darcey D, Tien LY, Vukmirovic AJ, Paula KY, et al. Posture-induced changes in intraocular, orbital, cranial, jugular vein, and arterial pressures in a porcine model. Investigative ophthalmology & visual science. 2023;64(15):22– 22.

63. Kim DS, Vaquer S, Mazzolai L, Roberts LN, Pavela J, Watanabe M, et al. The effect of microgravity on the human venous system and blood coagulation: a systematic review. Experimental Physiology. 2021;106(5):1149–1158.

64. Elahi MM, Witt AN, Pryzdial EL, McBeth PB. Thrombotic triad in microgravity. Thrombosis Research. 2024;233:82–87.

65. Lan M, Phillips SD, Archambault-Leger V, Chepko AB, Lu R, Anderson AP, et al. Proposed mechanism for reduced jugular vein flow in microgravity. Physiological Reports. 2021;9(8):e14782.

66. Williams ND, Mehlsen J, Tran HT, Olufsen MS. An Optimal Control Approach for Blood Pressure Regulation During Head-Up Tilt. Biological Cybernetics. 2019;113:149–159. doi:10.1007/s00422-018-0783-9.

67. Zhang B, Chen X, Qin W, Ge L, Zhang X, Ding G, et al. Enhancing cerebral arteriovenous malformation analysis: Development and application of patient-specific lumped parameter models based on 3D imaging data. Computers in Biology and Medicine. 2024;180:108977.

